# Establishing the baseline for using plankton as biosensor

**DOI:** 10.1101/795203

**Authors:** Vito P. Pastore, Thomas Zimmermann, Sujoy K. Biswas, Simone Bianco

## Abstract

Plankton is at the bottom of the food chain. Microscopic phytoplankton account for about 50% of all photosynthesis on Earth, corresponding to 50 billion tons of carbon each year, or about 125 billion tonnes of sugar[1]. Plankton is also the food for most species of fish, and therefore it represents the backbone of the aquatic environment. Thus, monitoring plankton is paramount to infer potential dangerous changes to the ecosystem. In this work we use a collection of plankton species extracted from a large dataset of images from the Woods Hole Oceanographic Institution (WHOI), to establish a basic set of morphological features for supporting the use of plankton as a biosensor. Using a perturbation detection approach, we show that it is possible to detect deviation from the average space of features for each species of plankton microorganisms, that we propose could be related to environmental threat or perturbations. Such an approach can open the way for the development of an automatic Artificial Intelligence (AI) based system for using plankton as biosensor.

## 1. INTRODUCTION

Plankton is a large class of aquatic organisms. Recent studies have proven that local and global environmental changes related to both manmade and natural events may profoundly disrupt the composition and dynamic of plankton populations[2]. Moreover, plankton of the species *Spirostomum ambiguum* has been used to detect the toxicity of volatile compounds through an analysis of morphological modifications due to the presence of an external perturbation[3]. Taken together, these micro- and macro-observations suggest that changes in plankton population, morphology and/or behavior over an established baseline of normal features, can be used a sensor of undesirable and unexpected events, even in the absence of a precise characterization of the external insult. In other words, plankton may indeed be used as a “canary in a coal mine”. For this purpose, a thorough characterization of the baseline of normal appearance or behavior needs to be carried out. In this paper, we introduce a computational methodology to build a reliable baseline of plankton morphology using features extracted from plankton images. Our methods may represent a first step in the development of a biosensor based on the analysis of significant deviation of plankton morphology.

## 2. MATERIALS AND METHODS

### 2.1 WHOI dataset

Images are extracted from a well-known database of plankton images, the Woods Hole Oceanographic Institution (WHOI) dataset (https://hdl.handle.net/10.1575/1912/7341), which comprises a collection of plankton images collected in situ by automated submersible imaging-in-flow cytometry with an instrument called Imaging FlowCytobot (IFCB). We extracted a selected set of 100 training images and 40 testing images for 10 species (see Fig 1), using the available datasets acquired during different years from 2006 to 2014.

**Fig 1.**
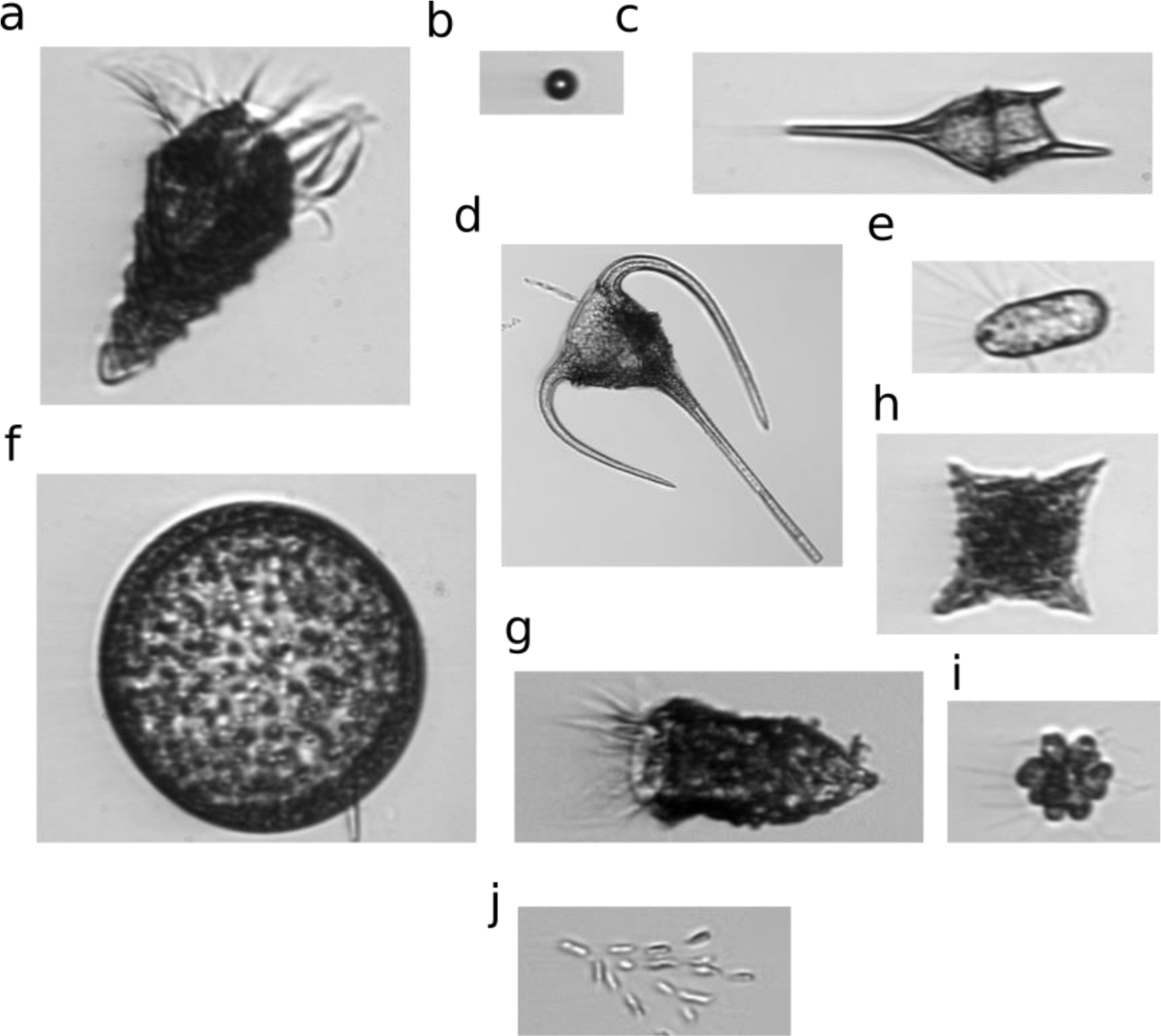
Collection of species extracted from the WHOI dataset. (a) laboea strobila, (b) baed, (c) ceratium humile, (d) ceratium tripos, (e) corethron, (f) coscinodiscus, (g) Tintinnid, (h) Dictyocha, (i) clusterflagellatae, (j) Dynobrion.

### 2.2 Features

We used OpenCV and python to process the plankton images extracted from the WHOI dataset to extract numerical image descriptors. The designed and implemented features can be divided into 4 characteristic classes, each of them describing different aspects of the image, embedding fundamental information for the purposes of classification and detection of perturbations.

#### 2.2.1 Segmentation

Images were segmented using OpenCV and python. A simple global bimodal segmentation based on Otsu algorithm [4] has shown to be effective for the extracted images. A closing morphological operation and a contour extraction provide both the contour and the cell body segmented image to be further processed for the features extraction.

#### 2.2.2 Moment-based features

Invariant moments are widely exploited in shape analysis and recognition, providing scaling, translation and rotation independent features. Moments-based features are not human interpretable as the geometric features are, however they have been shown to be efficient in encoding the morphological differences of plankton microorganisms. We used the normalized invariant moment proposed by Hu[5] (7 features) and the Zernike[6] moments up to order 5 (25 features).

#### 2.2.3 Texture-based features

A simple description of the texture is provided by the image’s gray levels patterns. Texture has been shown to be important in embedding morphological characteristics specific of the different plankton species [7], so we incorporated texture-based features into our set of image descriptors. We adopted both the Local Binary Pattern (LBP) descriptors (54 features) and the 13 Haralick descriptors. LBP describe the image’s texture providing information on the frequency of occurrence of the local gray levels pattern, codified as binary number. The Haralick features are extracted using the Gray Scale Co-occurrence Matrix (GSCM). The GSCM describes the distribution of gray levels counting the frequency of occurrence of each transition between couples of pixels.

We finally implemented 5 features extracted from the image’s gray-scale histogram. We extracted the mean value and the standard deviation of the histogram, that reflect globally how the gray levels are distributed into the image, as well as the kurtosis and the skewness of the peak and the entropy of the histogram. These measures provide information on the regularity and the texture patterns of the image.

#### 2.2.4 Geometric features

We used the extracted contours to compute a set of 14 simple shape descriptors. This set of features describes geometric properties of the contours, including area, eccentricity, rectangularity, aspect ratio, circularity, perimeter and convexity.

#### 2.2.5 Fourier Descriptors (FD)

We used the Fourier Descriptors (10 features) to encode global shapes features (low frequency FD). We computed the FD using the centroid distance. The centroid distance is a shape signature, a local representation for the contour of the image that has been showed to be resistant to noise and robust. The centroid distance is a 1-D function defined as:

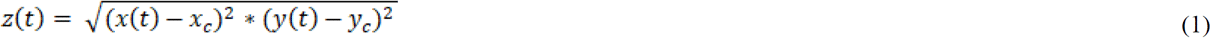

Where (x, y) are the contour points and (x_c_, y_c_) are the contour centroid’s coordinates.

### 2.3 One-class SVM

We exploited the one class SVM algorithm as described by Scholpoff [8]. Let us consider a set of observations (x_i_, y_i_= +1), where x_i_ is a m-dimensional real vector and y_i_= +1 refers to the data samples all belonging to a specific class. The one-class SVM is a classification algorithm learning a function taking +1 in the region of the features space where the majority of the training points fall, and -1 elsewhere. The algorithm maps the data into a high-dimensional space using a kernel. We adopted a Radial Basis Function (RBF) as a kernel, for the plankton images analysis described in this manuscript. The one-class SVM can be interpreted as a two-class SVM aimed in learning the hyperplane dividing the class represented by the training data from the origin, which is the only member of a second class. Such a hyperplane can be defined as:

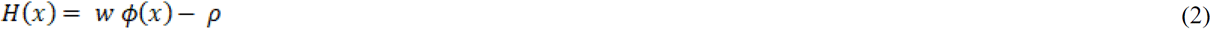

The one-class SVM’s training maxims the distance of the hyperplane from the origin, i.e., it maxims the quantity 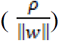 solving a quadratic problem. The hyperplane is further used as a classification rule to assess if a test sample either belongs to the trained class.

## 3. RESULTS

### 3.1 Features importance in classification

We used a random tree classifier-based approach to determine a criterion of importance of the designed features in classifying the extracted plankton species. Fig 2a shows the results obtained using the python implementation provided by the scikit-learn library [9]. Such an implementation adopts a Gini Importance or Mean Decrease in Impurity (MDI) metrics in calculating each feature importance [10]. Both the shape-based and the textures-based features result important for separating the analyzed species (cf. Fig 2a). As expected, the geometric features show the highest coefficient of importance (i.e., they are the most exploited in splitting the nodes of the decision trees). In the texture-based features, the Haralick features are the most important, proving the distribution and pattern of the gray values are much exploited by the algorithm in classifying the analyzed plankton species.

**Fig 2.**
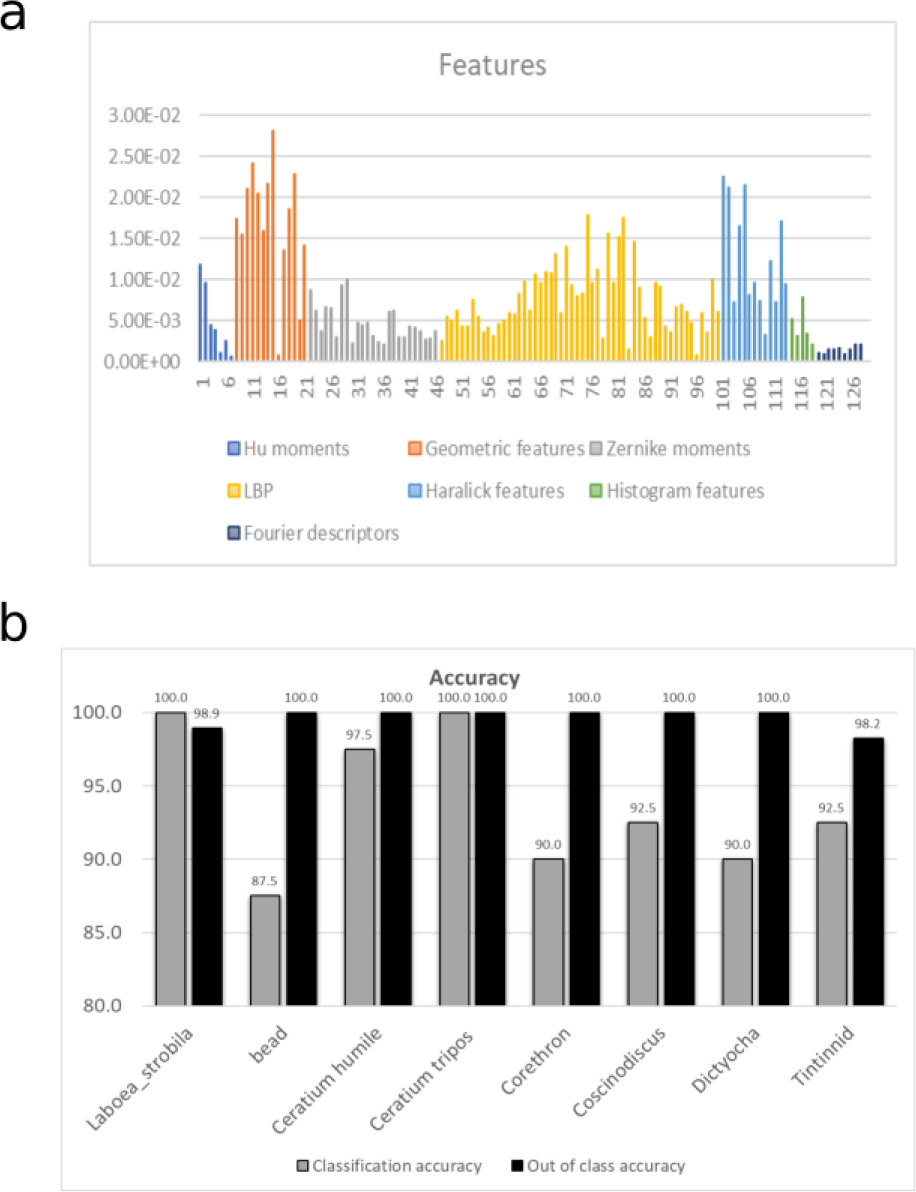
(a) Features score of importance provided by the decision tree algorithm for the 10 species included into the analysis, (b) One-class SVM testing accuracy for the different training species for both in-class and out of class samples.

### 3.2 One-class SVM detector performances

We trained a one-class SVM detector using the 100 training images extracted for each of the species selected from the WHOI dataset. We further tested each of the detectors using the 40 images testing images of the training class as positive samples (i.e., samples belonging to the class) and the remaining 280 testing images, corresponding to the other class into the dataset, as negative samples (i.e., samples not belonging to the class). We trained a one-class SVM detector for each of the first eight species listed in Fig 1. The training procedure is supervised, i.e., we used the plankton species annotated labels during training. Fig 2B shows the obtained performances. The trained detectors reveal a very high testing classification accuracy, equal to (93.8 ± 4.8) % and an almost ideal out of class detection testing accuracy, reaching a value of (99.6 ± 0.7) %. The obtained results confirm the high accuracy of the designed features in separating the extracted plankton species.

In order to test if our computational methodology is able to produce a reliable baseline to infer the presence of morphological modifications, we used images of two additional species (*Dynobrion* and *clusterflagellatae*) to test the system composed of all the trained detector. Fig 3 shows a schematic description of the pipeline designed to perform such an analysis. We used the 140 images for each of the two species to test if all the implemented detectors globally recognize them as out of class. As Fig 4 shows, the *Dynobrion* is locally detected as out of class with an average accuracy equal to (99.9 ± 0.3) %, while the *clusterflagellatae* is locally recognized as out of class with an average accuracy of (99.3 ± 1.3) %. For each of the 140 samples correspondent to these two classes, we computed the global response of all the trained detectors: the sample is defined as global out of class only if all the other detectors discard it. The resulting absolute number of global out of classes samples corresponds to 139 for the *Dynobrion* and 132 for the *clusterflagellatae*.

**Fig 3.**
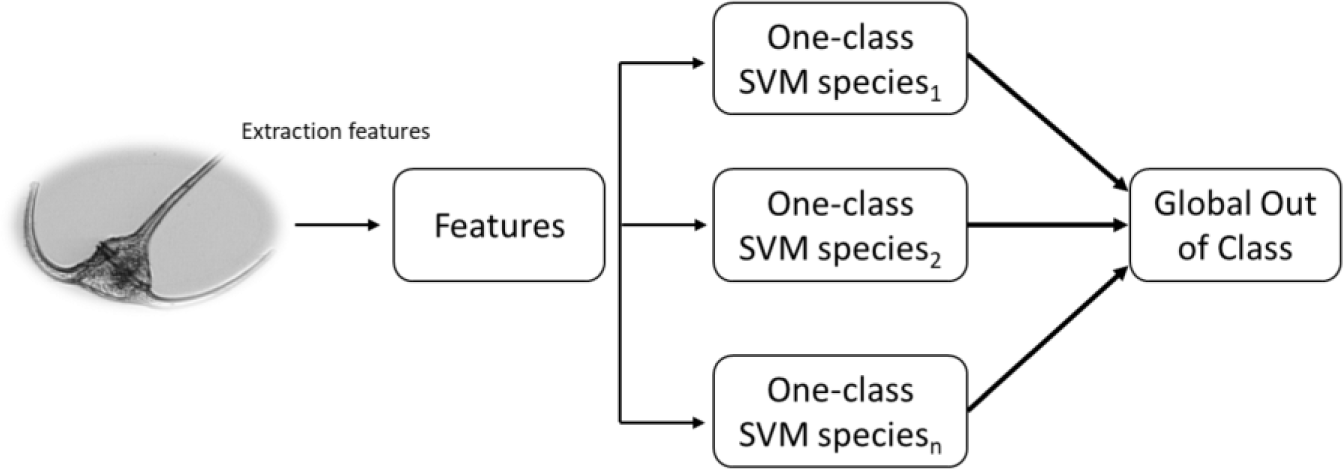
Schematic description of the pipeline designed for the testing our computational methodology. All the trained detectors are inputted with the multi-dimensional samples resulting from the feature extraction process. The sample is defined as global out of class if all the one-class SVMs respond with a -1 to the sample, that is, all the detectors recognize it as external to the training class.

**Fig 4.**
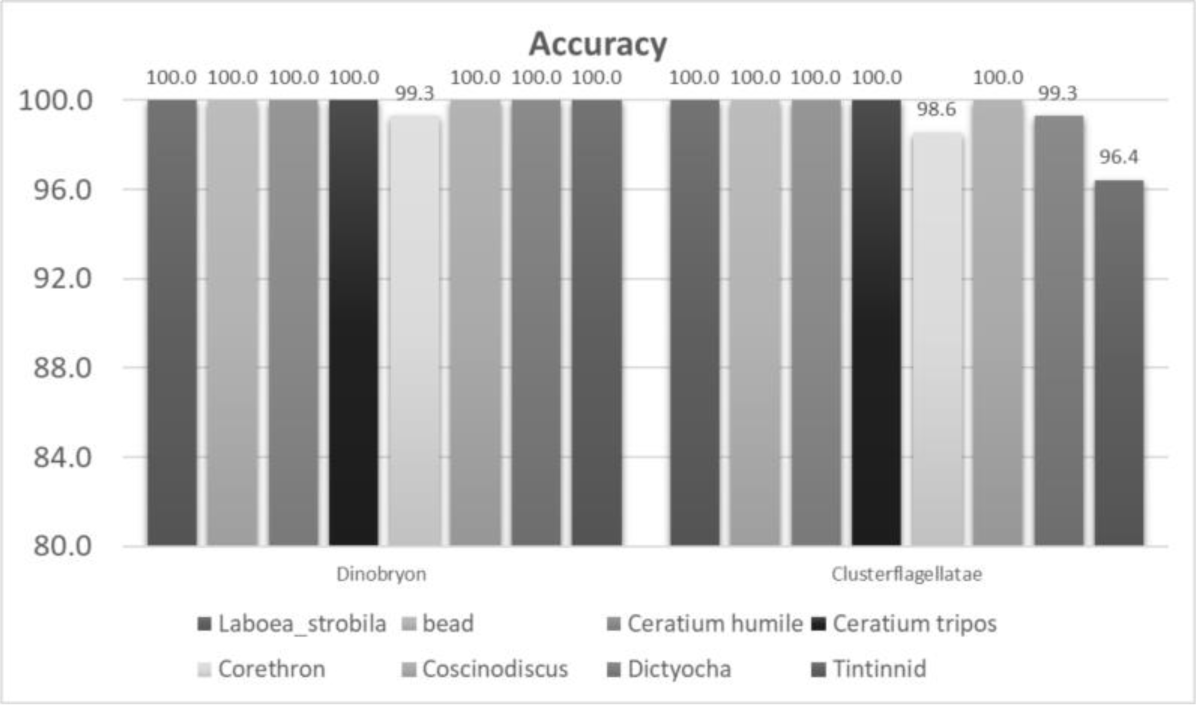
One-class SVM accuracy for the different training species in detecting *Dinobryon* and *Clusterflagellatae* as out of class.

## 4. DISCUSSION

In this manuscript we extracted 128 morphological features describing different aspects of plankton images from the WHOI dataset. We defined a binary out of class detector based on the one-class SVM. We showed that such approach can reach a testing accuracy of about 94% on the collection of species extracted from the WHOI dataset. Such result confirms the efficiency of the designed features in embedding the plankton morphological information and separating the different species. We then used 2 of the extracted species as benchmark to evaluate the global response of the detectors. As showed in the results, the detectors globally recognized the two species as out of class with an average accuracy higher than 95%. The aim of this work is to establish a baseline of plankton morphological descriptors to use plankton as biosensor. The system developed with a basic one class-SVM based algorithm is able to recognize, as global out of class, two species selected from the dataset and excluded from the detectors training. This suggests that, if plankton morphology is a proxy for the presence of environmental perturbations, then a system of algorithms based on the work we propose could be used to reveal such changes. This approach may lead to the development of more complete method to use plankton as biosensor.

## Acknowledgment

This material is based upon work supported by the National Science Foundation under Grant No. DBI-1548297. Disclaimer: Any opinions, findings and conclusions or recommendations expressed in this material are those of the authors and do not necessarily reflect the views of the National Science Foundation.

